# Combinatorial design of chemical-dependent protein switches for controlling intracellular electron transfer

**DOI:** 10.1101/645978

**Authors:** Bingyan Wu, Joshua T. Atkinson, Dimithree Kahanda, George. N. Bennett, Jonathan J. Silberg

**Affiliations:** Biochemistry & Cell Biology Graduate Program, Rice University, MS-140, 6100 Main Street, Houston, TX, 77005; Department of Biosciences, Rice University, MS-140, 6100 Main Street, Houston, TX, 77005; Systems, Synthetic, & Physical Biology Graduate Program, Rice University, MS-180, 6100 Main Street, Houston, TX, 77005; Department of Chemical & Biomolecular Engineering, Rice University, 6100 Main Street, MS-362, Houston, Texas 77005; Department of Bioengineering, Rice University, 6100 Main Street, MS-142, Houston, Texas 77005

**Author notes:** To whom correspondence should be addressed: Jonathan J. Silberg, Biosciences Department, 6100 Main Street, Houston TX 77005.

**Keywords:** allostery, domain insertion, electron transfer, ferredoxin, growth selection

## Abstract

One challenge with controlling electron flow in cells is the lack of biomolecules that directly couple the sensing of environmental conditions to electron transfer efficiency. To overcome this protein component limitation, we randomly inserted the ligand binding domain (LBD) from the human estrogen receptor (ER) into a thermostable 2Fe-2S ferredoxin (Fd) from *Mastigocladus laminosus* and used a bacterial selection to identify Fd-LBD fusion proteins that support electron transfer from a Fd-NADP reductase (FNR) to a Fd-dependent sulfite reductase (SIR). Mapping LBD insertion sites onto structure revealed that Fd tolerates domain insertion adjacent to or within the tetracysteine motif that coordinates the 2Fe-2S metallocluster. With both classes of the fusion proteins, cellular ET was enhanced by the ER antagonist 4-hydroxytamoxifen. In addition, one of Fds arising from ER-LBD insertion within the tetracysteine motif acquires an oxygen-tolerant 2Fe-2S cluster, suggesting that ET is regulated through post-translational ligand binding.

## INTRODUCTION

In redox synthetic biology, non-native electron transfer (ET) pathways are constructed to control electron flow within cells and between cells and extracellular materials^1–3^. Within the cytosol, synthetic control over ET can aid in optimizing the flux through metabolite pathways by altering the cofactor specificity of oxidoreductases^4–6^, increasing electron transfer (ET) efficiency between pairs of proteins through fusion^7, 8^, and generating synthetic cells that insulate ET pathways from off-pathway reactions^9–11^. Regulating ET is also critical for developing the next generation of bioelectronics and biosensors^12^. Recent studies have shown that diverse bacteria can couple their metabolism with extracellular materials^13–15^. Photosensitive extracellular materials have been used to create light-driven enzymatic synthesis through direct coupling with oxidoreductases or by precipitating these materials on cells to create shells that harvest light for cell metabolism^16–18^. Additionally, proteins that mediate extracellular ET have been transplanted into non-native hosts to enable synthetic electrical communication between cells and materials^19, 20^. While hybrid cells and materials have been reported that leverage synthetic biology for *in situ* sensing and chemical synthesis, these studies typically control ET at the transcriptional level, limiting the speed at which ET can be regulated for different applications.

One challenge with controlling ET within cells is our limited understanding of the mechanisms by which this process is regulated in nature. A wide range of protein electron carriers (PECs) have been described that control ET^21^. In the low potential range, ferredoxins (Fds) and flavodoxins (Flds) are frequently used to regulate ET^22, 23^. Many organisms contain multiple low potential PECs, with some organisms containing >50 Fds and Flds^24^. Biochemical and proteomic studies have provided evidence that PEC paralogs can differ in their specificity for their partner oxidoreductases^25, 26^, allowing them to function as ET hubs, directing different proportions of electrons between the coexpressed partner proteins. Cells can gain additional control over ET by differentially regulating the gene expression of PEC paralogs^27, 28^. Recently, evidence emerged that Fds may at times use post-translational control to regulate ET in cells. Nitrogenase-protecting Fds use oxidation to alter their conformation and binding to nitrogenase as a mechanism to protect their partner protein from oxidative damage^29, 30^. Additionally, phosphorylated and calcium-bound Fds have been reported, suggesting that fast regulation of ET has arisen during evolution^31, 32^. However, very few natural proteins have been discovered that use post-translational mechanisms to control ET. This limits our ability to program ET for metabolic engineering and bioelectronic applications.

Recently, a protein electrical switch was engineered through rational design that exhibits chemical-dependent ET^3^. In this study, Fd-family sequence information was used to identify backbone locations in a thermostable 2Fe-2S Fd from *Mastigocladus laminosus* (*Ml*-Fd) that tolerate fission without losing the ability to support ET in cells^33^. All four of the rationally-designed split Fds (sFds) were unable to support ET transfer unless they were fused to pairs of peptides that stabilized their interaction^3^. Additionally, the fragments from one variant could be fused to the ends of a ligand-binding domain (LBD) to create a chemical-dependent protein switch^3^. This topological mutant arose from insertion of the human estrogen receptor (ER) LBD after Gly35 in *Ml-*Fd, a location that is outside of the tetracysteine motif that coordinates the 2Fe-2S metallocluster^33^. While this study provided evidence that domain insertion can be used to create electrical protein switches, using an approach similar to that described for creating fluorescent biosensors for estrogenic compounds^34^, it only examined one of the possible designs for a Fd containing a ER LBD.

To better understand where Fds tolerate insertion of LBDs, we used transposon mutagenesis to randomly insert the ER LBD at different locations within *Ml*-Fd^35–37^, and we subjected this library to a cellular selection for Fd ET. We identified three novel topological mutants that support cellular ET. These proteins all presented enhanced ET in cells when exposed to the ER antagonist 4-hydroxytamoxifen (4-HT). Purification of one variant arising from ER-LBD insertion within the *Ml*-Fd tetracysteine motif revealed that this protein acquires an oxygen-tolerant 2Fe-2S cluster in the absence of 4-HT, suggesting that some of these proteins regulate ET through chemical-induced changes in the holoprotein structure.

## Materials and Methods

### Materials

DNA oligos were obtained from Integrated DNA Technologies. Phusion DNA polymerase was purchased from Thermo Fisher. Restriction enzymes and T4 DNA ligase were purchased from New England Biolabs. Chloramphenicol, kanamycin, isopropyl β-D-1-thiogalactopyranoside (IPTG), and dithiothreitol (DTT) were purchased from Research Products International. Anhydrotetracycline (aTc) and 4-hydroxytamoxifen (4-HT) were purchased from Sigma-Aldrich.

### Transposon Design

To create a transpon for inserting unique BsaI sites at different locations within the Fd gene, we PCR amplified the Entranceposon M1-CamR (F760, Thermo Fisher Scientific) using primers that modify the R1 sequences to encode BsaI restriction sites. The amplicon was cloned into pUC19 using HindIII and XbaI, and the chloramphenicol resistance cassette (*chl^R^*) was replaced with a kanamycin resistance cassette (*kan^R^*) using Gibson Assembly^38^. The transposon-propagation plasmid (pUC19-Mu-BsaI-Kan^R^) encodes a transposon (Mu-BsaI-Kan^R^) that is similar in sequence to a previously described transposon^48^. In our transposon, the 4 base pair single-stranded overhangs generated by BsaI digestion (CTGC and GCTG) are distinct, since they were designed to be compatible with the N-terminal linker sequence in our synthetic gene encoding the ER-LBD, which encoded an Ala-Ala dipeptide at the beginning. In addition, our transposon contains *kan^R^* rather than *chl^R^*. To generate Mu-BsaI-Kan^R^ for MuA reactions, it was excised from transposon-propagation plasmid using HindIII and BglII.

### Staging Library

The gene encoding *Mastigocladus laminosus* (*Ml*-Fd) was PCR amplified to create an amplicon encoding the *Ml*-Fd gene flanked by BbsI restriction sites. This DNA was cloned into pFd007, which uses an aTc-inducible promoter for expression^3^, and the resulting *chl^R^* vector (pFd007-staging) was used for the transposition reaction. Transposition was performed in a 20 µL reaction by mixing MuA reaction buffer, 100 ng linear Mu-BsaI-Kan^R^, 500 ng pFd007-staging, and 0.22 μg MuA (Thermo Fisher Scientific, F750). The reaction was incubated at 30 °C for 16 hours and then 75 °C for 10 minutes to heat inactivate MuA. The reaction product was purified using a DNA Clean and Concentrator-5 kit (Zymo Research), and the entire sample was transformed into electrocompetent *E. coli* XL1-Blue. Cells were allowed to recover in SOC medium at 37 °C for 1 hour prior to plating onto LB-agar plates containing 34 μg/mL chloramphenicol and 50 μg/mL kanamycin. This antibiotic combination selects for the antibiotic resistance cassettes in pFd007-staging (*chl^R^*) and Mu-BsaI-Kan^R^. Approximately 25,000 cfu were obtained. Assuming that the cells doubled one time during the SOC recovery, this indicates that our library contains 12,500 unique vectors. To generate the staging library, cells were harvested from plates by adding LB, scraping colonies into the LB, and pooling the cell slurries from each plate. The vector ensemble was purified from the cell slurry using a Miniprep Kit (Qiagen).

### Size-Selected Library

The staging library contains vectors with transposons inserted in the vector and in the gene of interest^39^. To isolate Fd genes containing a transposon, the staging library was digested with BbsI and size selected using a 0.8% agarose gel. Linear DNA corresponding to a Fd gene containing a single transposon (1,700 bp) was excised from the gel and purified using a Gel DNA recovery kit (Zymo Research). The Fd-transposon gene fusions were cloned into pFd007-staging using Golden Gate Assembly^40^. The product of the reaction was purified using a DNA Clean and Concentrator kit (Zymo Research), the entire product was transformed into electrocompetent *E. coli* XL1-Blue, cells were allowed to recover in 1 mL of SOC at 37 °C for 1 hour, and cells were plated onto LB agar plates containing 34 μg/mL chloramphenicol and 50 μg/mL kanamycin. The colonies obtained after overnight growth (∼13,000) were harvested from plates, and the resulting library was purified.

### Domain-Insertion Library

The gene encoding the ER-LBD (residues 302-552) flanked by linkers (AAAGGGGSGGGGS at the N-terminus, GGGGSGGGGSAAA at the C-terminus) was PCR amplified from a previously described template using primers that add BsaI restrictions sites at the ends of the amplicon^3^. The size-selected library was digested with BsaI, the vector (∼3,500 bp) was separated from the transposon using using gel electrophoresis, and the vector DNA was purified using a Zymo clean gel DNA recovery kit (Zymo Research). This purified DNA and ER-LBD gene were assembled using Golden Gate Assembly^40^. The product of this reaction was purified using a DNA Clean and Concentrator kit (Zymo Research) and transformed into *E. coli* XL1-Blue using electroporation. Following recovery in SOC medium at 37 °C for 1 hour, cells were plated onto LB agar containing 34 μg/mL chloramphenicol. Following a 16 hour incubation at 37°C, ∼1,000 cfu were obtained. These colonies were harvested and pooled. The final domain-insertion library was obtained by purifying plasmids from this cell slurry using a QIAprep Spin Miniprep Kit (Qiagen, Inc.).

### Library Selection

*E. coli* EW11, a strain lacking a functional sulfite reductase gene^11^, was transformed using electroporation with pSAC01 (100 ng), a plasmid that constitutively expresses *Zea mays* FNR and SIR^3^, and the domain-insertion library (100 ng). Cells were recovered in SOC at 37 °C for 1 hour, pelleted by centrifugation, and resuspended in M9sa medium^3^, which contains sulfate as the only sulfur source. Resuspended cells were spread onto four types of agar plates, including plates containing M9sa, M9c, which is made by adding methionine and cysteine to M9sa, M9sa containing 200 ng/mL aTc, and M9sa containing 200 ng/mL aTc and 50 μM 4-HT. All plates contained 34 μg/mL of chloramphenicol and 100 μg/mL of streptomycin to select for library vectors and pSAC01, respectively. Plates were incubated at 37 °C for 72 hours prior to assessing growth. The topological mutants obtained from the selection are made up of a sFd whose fragments are fused to the ER LBD termini. For this reason, we name each topological mutant using the residue after which the ER LBD was inserted, *e.g.*, sFd-25-ER.

### Complementation Analysis

Electroporation was used to transform *E. coli* EW11 with pSAC01 and a plasmid that expresses either *Ml*-Fd^3^, a C42A mutant of *Ml-*Fd that lacks a cysteine required to coordinate a 2Fe-2S cluster^3^, or a topological Fd mutant. Single colonies containing both plasmids were obtained by growing cells on LB-agar plates containing 34 μg/mL of chloramphenicol and 100 μg/mL of streptomycin. Individual colonies obtained from overnight incubations were used to inoculate liquid cultures (100 μL) containing M9c with 0.5% DMSO, 34 μg/mL of chloramphenicol, and 100 μg/mL of streptomycin. These cells were grown for 16 hours while shaking in an Infinite m1000 Pro plate reader (Tecan) at 37 °C with shaking at 217 rpm and an amplitude of 1.5 mm in double-orbital mode. An aliquot (2 μL) of each starting culture was diluted 1:50 in fresh M9sa medium (100 µL) in a flat bottom Costar 96-well plate (#3595, Corning). These cultures were then grown in the presence and absence of aTc (200 ng/mL) and 4-HT in an Infinite m1000 Pro plate reader (Tecan) at 37 °C with shaking at 217 rpm and an amplitude of 1.5 mm in double-orbital mode. Cell growth was assessed every 10 minutes by reading optical density (OD) at 600 nm for 48 hours. To minimize evaporation, the 96-well plate was wrapped with parafilm. Control experiments (*Ml*-Fd and C42A) were performed in wells that were separated from each by at least one row to avoid cross contamination.

### Protein Purification

*Ml*-Fd was purified as previously described^21^. The gene encoding sFd-55-ER was PCR amplified and cloned into pET28b using Golden Gate Assembly^40^. *E. coli* Rosetta(DE3) transformed with this plasmid were grown at 37 °C in 4 L of LB containing 50 μg/mL kanamycin and 100 μM ferrous ammonium sulfate to mid log phase (OD ∼0.7). Protein expression was induced by adding 0.1 mM IPTG. Following induction, cells were grown overnight at 25 °C while shaking at 250 rpm, harvested by centrifugation (4,000 xg) and resuspended in lysis buffer, which contained 50 mM Tris pH 8, 5 mM dithiothreitol (DTT), and cOmplete Mini EDTA-free protease inhibitor tablet (Roche Diagnostics GmbH). Resuspended cells were split into 30 mL fractions, sonicated on ice with a Qsonica sonicator, 4435 Q55 Sonicator Microprobe (1/4“), 70% amplitude, 1s on 1s off pulse for 8 minutes twice. Clarified lysates were generated by centrifugation (20,000 xg) and loaded onto a DE52 anion exchange column (Whatman) that had been equilibrated with TED buffer (50 mM Tris pH 8, 1 mM EDTA, and 5 mM DTT). The column was washed with TED, and TED containing 50, 100, 150, 200, 250, 300 and 500 mM NaCl was used to elute protein. The brown eluent, which eluted at 250 mM NaCl, showed a band of expected size (41 kDa) on an SDS-PAGE. This fraction was diluted five times with TED buffer and loaded onto HiTrap Q XL column using an AKTA Start FPLC system (GE Healthcare). The column was washed with TED buffer, and a linear NaCl gradient in TED was used to elute the protein. Those fractions appearing brown were analyzed for purity using SDS-PAGE. All fractions contained a major band having the expected molecular weight (41 kDa). Those fractions appearing homogeneous were pooled and concentrated using Amicon Ultra 10k MWCO spin column. This protein was further purified using a size exclusion column (HiLoad16/600 Superdex 200pg, GE Healthcare) equilibrated with TED containing 100 mM NaCl. Protein appearing homogeneous by SDS-PAGE was pooled and used for all subsequent analysis.

### Whole Cell Fluorescence

The RFP gene was PCR amplified and cloned using Golden Gate Assembly into aTc-inducible vectors that express *Ml*-Fd (pFd007), sFd-25-ER (pFd-ER-25), sFd-30-ER (pFd-ER-30), sFd-35-ER (pFd-ER-35), sFd-54-ER (pFd-ER-54) and sFd-55-ER (pFd-ER-55). This yielded vectors that express Fds with RFP fused to their C-terminus through 12 amino acid linker (GGSGGSGGSGGS). Each vector was transformed into *E. coli* EW11 with pSAC01, cells were grown on LB-agar plates containing 34 μg/mL of chloramphenicol and 100 μg/mL of streptomycin, and individual colonies were used to inoculate liquid cultures (100 μL) containing M9c, 0.5% DMSO, 34 μg/mL of chloramphenicol, and 100 μg/mL of streptomycin. These cells were grown for 16 hours in a Spark plate reader (Tecan) at 37 °C with shaking at 90 rpm at an amplitude of 3 mm in double-orbital mode. An aliquot (2 μL) of each starting culture was diluted 1:50 in fresh M9c medium (100 µL) in a Nunc™ Edge 2.0 96-Well Plates (Thermo Fisher). These cultures were grown in the presence and absence of aTc (200 ng/mL) and 4-HT in a Spark plate reader (Tecan) at 37 °C using a similar procedure. Cell growth was assessed every 5 minutes by reading optical density (OD) at 600 nm for 24 hours as well as whole cell red fluorescence (λ_ex_ = 560 nm, λ_em_ = 650 nm). The ratio of emission to OD was calculated for stationary phase cultures. To minimize evaporation, 96-well plates were wrapped with parafilm, and grown in a Spark humidity cassette.

### Spectroscopic Characterization

Purified sFd-55-ER was dialyzed into TED buffer containing 100 mM NaCl. Protein concentration was determined spectrophotometrically using the calculated extinction coefficient (ℇ_280nm_ = 32,890 M^-1^cm^-1^). A Varian Cary 50 spectrophotometer and a JASCO J-815 spectropolarimeter were used to obtain absorbance and circular dichroism spectra, respectively. Samples contained 40 µM sFd-55-ER in a cuvette having a 1 cm path. For comparison, absorbance and circular dichroism spectrums were obtained for 40 μM *Ml-*Fd (ℇ_280nm_ = 9,190 M^-1^cm^-1^).

### Midpoint potentials

Electrochemical measurements were carried out anaerobically using a DropSens Stat 8000P Potentiostat. Glass vial with a three-electrode system was continuously purged with Ar until the time of the electrochemical measurement, which used a 1 M, KCl Ag/AgCl electrode as the reference electrode and a platinum wire as the counter electrode. Edge-plane graphite electrode (EPG) was used as the working electrode for thin film electrochemistry and potentials were reported relative to the standard Hydrogen electrode (SHE). Electrodes were cleaned and placed in the glass cell containing 23.5°C mixed buffer solution (5 mM sodium acetate/MES/MOPS/TAPS/CHES/CAPS) pH 8.0 containing 100 mM NaCl. Aliquot of 431 μM *Ml*-Fd (7 µL) or 302 μM sFd-ER-55 (10 µL) were applied directly on the PGE electrode surface with or without 1 μL of 50 mM 4-HT, which was saturated in TED buffer. Thin film was incubated for 5 minutes, while blowing the surface with Ar gas. Square wave voltammograms were collected at 23.5°C at 10 Hz, immediately following placement of the PGE electrode in the glass cell containing buffer solution. Electrochemical signals were analyzed using QSOAS open-source software. Two experiments were performed on each protein with or without 4-HT, under identical conditions.

### Statistics

The results from growth complementation and whole cell fluorescence measurements represent the mean and standard deviation of biologically independent samples (n = 3). All reported *P* values were obtained using two-tailed, independent t-tests.

## RESULTS AND DISCUSSION

### Library construction

The pioneering studies by Frances Arnold and coworkers revealed that artificial proteins can be created in the lab by mimicking evolution^41^ In this directed evolution approach^42^, proteins with novel functions are discovered by creating combinatorial libraries of mutants and mining those libraries for biomolecules with the desired functional properties using a selection or a screen. In cases where large fitness jumps are desired, libraries encoding topological mutants of proteins are increasingly used, such as random domain insertion libraries for creating protein switches^43, 44^. Several library approaches have been described for generating domain insertion libraries, including nuclease-based and transposase-mediated methods^35, 36, 45, 46^. Among these approaches, transposase mutagenesis is appealing to use for creating protein switches because it avoids deletions that can arise with nuclease-based methods^37^. Transposase mutagenesis has recently been used to discover chemical-dependent protein switches by targeting lactamase^35^, CRISPR-Cas9^47^, GFP^48^, and an amino-acyl tRNA synthetase^36^ for random domain insertion.

To test whether transposase-mediated protein engineering can be used to discover novel sFd-ER that support ET in cells, transposase-mutagenesis was used to generate a library of sFd-ER (Figure 1a). We first created a synthetic transposon (Figure 1b) that contains a selectable marker (*kan^R^*) and transposase-binding sites (R1R2/R2R1) that are flanked by BsaI restriction site. This DNA is similar to a recently described synthetic transposon^48^, with one minor difference. The single stranded DNA overhangs adjacent to the transposase binding sites (R1R2 and R2R1), which encode a portion of the linkers that ultimately connect the ER-LBD and Fd, were modified so that they encode an Ala-Ala dipeptide. A three-step protocol was then used to create the vector library. In this protocol, MuA was initially used to randomly insert the synthetic transposon into a vector containing the *Ml*-Fd gene. To obtain the staging library, the product of this reaction was transformed in *E. coli* XL1-Blue, and vectors containing an inserted transposon were selected by growing cells on LB-agar plates containing kanamycin and chloramphenicol. This selection yielded ∼12,500 unique colonies, which exceeds the number of possible variants in our library (n = 6,482). The Fd-transposon gene hybrids were excised from this ensemble of vectors and subcloned into a vector that uses TetR to control Fd gene expression^3^. This size-selected library was transformed into *E. coli* XL1-Blue, and cells were grown on LB-agar plates to select for the desired vectors. This cloning step yielded ∼6,500 unique colonies, which exceeds the number of Fd-transposon gene hybrids (n = 296 unique insertion sites) that our method can create. To generate the final domain insertion library, we subcloned the gene encoding the ER-LBD (residues 302-552) in place of the synthetic transposon. This final step cloning yielded more than 500 unique colonies. In cases where this gene is inserted in frame, it is fused to the Fd through flexible linkers at its N- and C-termini (AAAGGGGSGGGGS and GGGGSGGGGSAAA, respectively).

**Figure 1.**
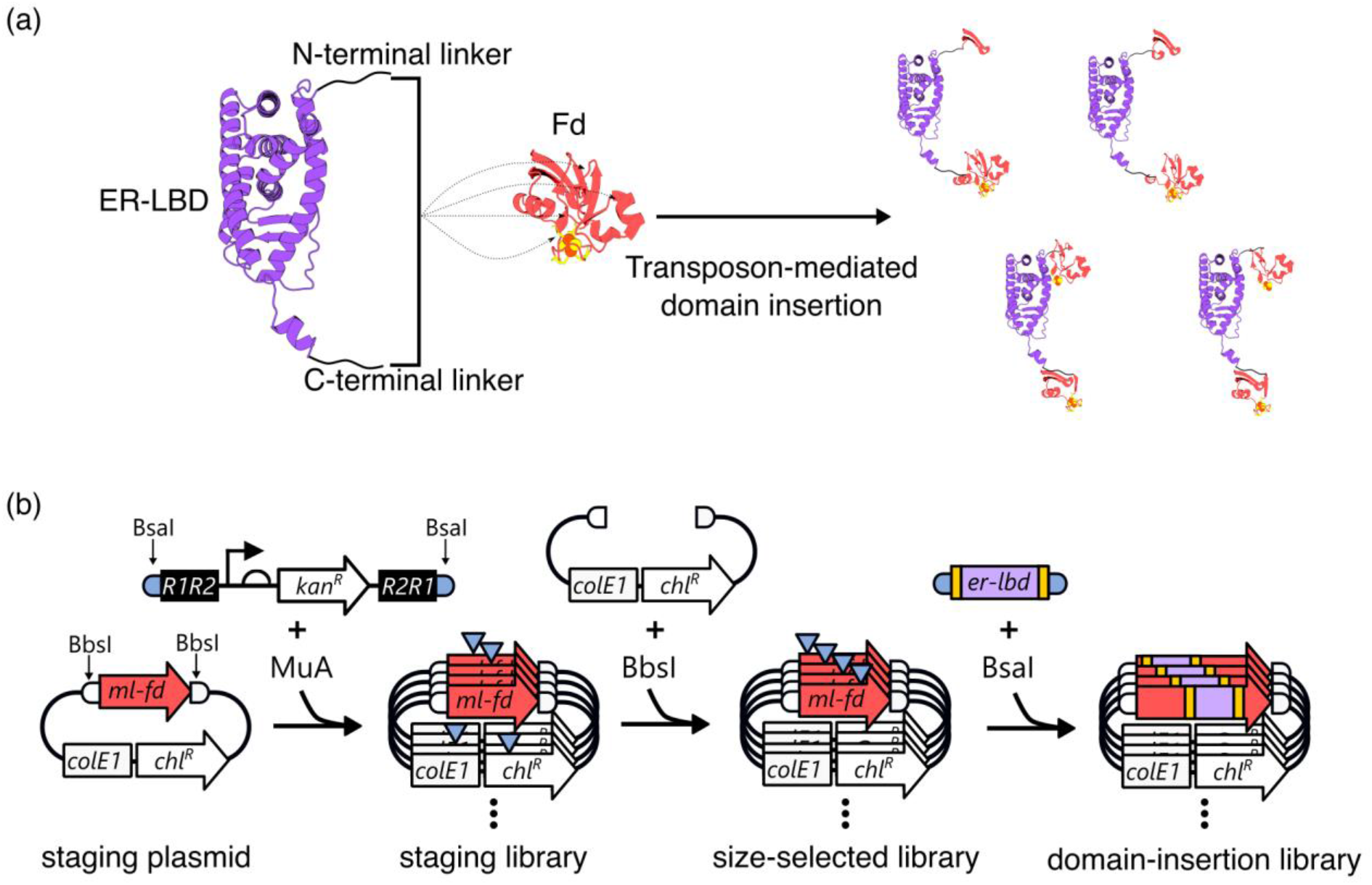
Library creation using random domain insertion. (**a**) Transposon-mediated domain insertion of the ER-LBD into *Ml*-Fd generates a library of topological mutants. (**b**) Three-step protocol for library generation. In step 1, MuA inserts a synthetic transposon at different position of the staging plasmid. Restriction digestion of the resulting staging library with BbsI yields *Ml*-Fd containing and lacking a transposon insertion. In step 2, the *Ml*-Fd-transposon gene hybrids are isolated and cloned into a vector to create the size-selected library. In step 3, the transposon is replaced with a gene encoding ER-LBD to generate the domain-insertion library.

### Selecting for protein electron carriers

To identify sFd-ER that coordinate a 2Fe-2S cluster and are able to support ET from a donor to an acceptor protein, we transformed the domain insertion library into *E. coli* EW11^11^, a strain with a chromosomal mutation within the gene (*cysI*) that encodes sulfite reductase. Because this strain is unable to grow on minimal medium containing sulfate as a sulfur source^11^, we transformed our library of vectors into cells along with a vector that constitutively express *Zea mays* FNR and SIR^3^. We then selected for growth on agar plates containing sulfate as the only sulfur source, aTc to induce sFd-ER expression, and 4-HT. This latter chemical was included in our growth medium because structural studies have shown that this and other antagonists stabilize an ER-LBD conformation with the N- and C-termini in closer proximity than that observed in the absence of bound ligand or in the presence of bound agonists^49–51^. With this selection, growth complementation was expected to only occur in cases where a sFd-ER was able to support ET from FNR to SIR (Figure 2a).

**Figure 2.**
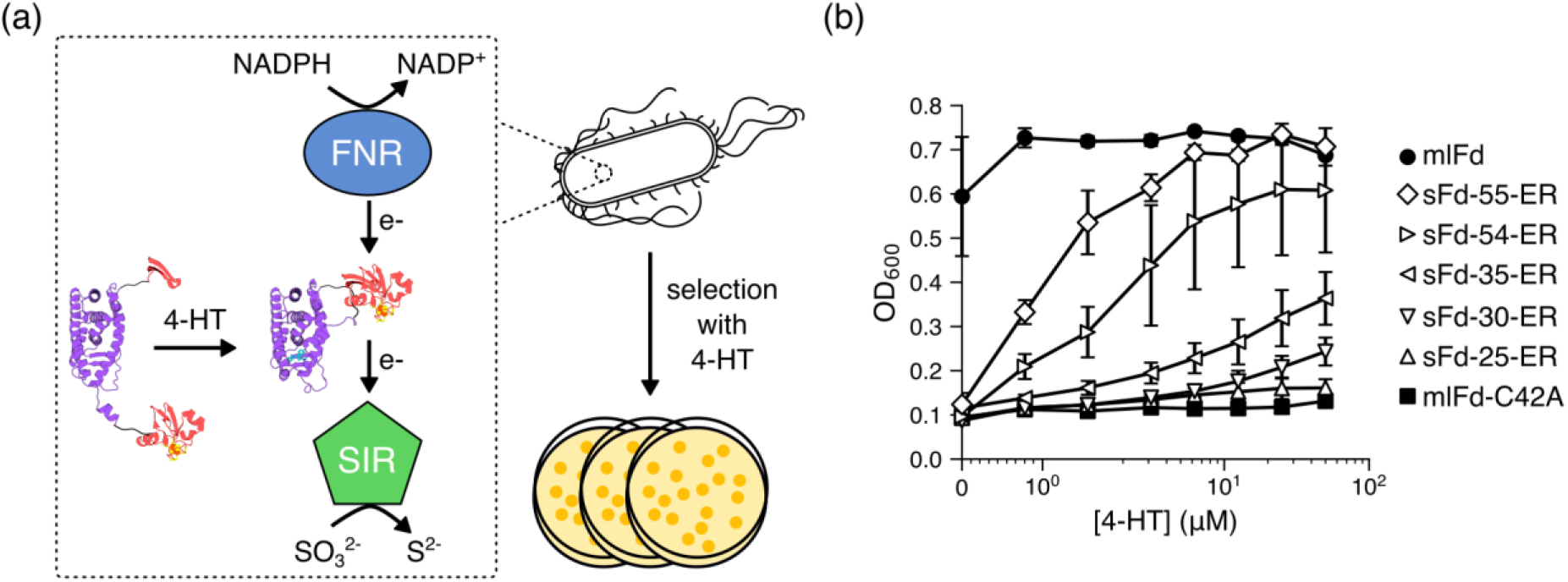
Selecting for ferredoxins that exhibit electron transfer. (**a**) Cells are only able to grow on minimal medium containing sulfate as the sulfur source if they express a Fd that supports ET from FNR to SIR. Selection of the domain-insertion library in presence of 4-HT at 37°C yielded five topological mutants, including ER-LBD insertions after *Ml*-Fd residues Gln25, Ala30, Gly35, Leu54, and Ile55. (**b**) Growth complementation observed after 48 hours with *E. coli* EW11 expressing FNR, SIR and the different Fd topological mutants in the presence of different 4-HT concentrations. After a 48 hour aerobic incubation at 37°C, absorbance at 600 nm was measured to quantify complementation. As a frame of reference, experiments were performed using *Ml*-Fd and *Ml*-Fd-C42A. Error bars represent ±1σ calculated from 3 biologically independent samples. Compared to the C42A mutant, growth was significantly enhanced in *Ml*-Fd under all conditions (*P* = 6.38×10^-3^), in sFd-55-ER, sFd-54-ER, and sFd-35-ER in the presence of ≥ 0.781 µM 4-HT (*P* = 3.31 × 10^-4^, 8.32 × 10^-3^, 8.68 × 10^-3^, respectively), in sFd-30-ER in the presence of ≥ 25 µM 4-HT (*P* = 8.85 × 10^-3^). sFd-25-ER did not show enhanced growth under any condition (*P* > 0.05).

Selection of our domain-insertion library on medium containing sulfate as the only sulfur source yielded 36 colonies that complemented growth in the presence of 4-HT, which represented ∼3% of the colonies plated under these conditions. Growth complementation performed in the absence of 4-HT yielded a smaller percentage of colonies. Sequencing colonies obtained from this selection (+4-HT) identified 5 unique topological mutants, which arose from insertion of the ER-LBD within *Ml*-Fd following residues 25 (sFd-25-ER), 30 (sFd-30-ER), 35 (sFd-35-ER), 54 (sFd-54-ER), and 55 (sFd-55-ER). These variants all arose from in frame insertions. Terminal fusions (LBD-Fd and Fd-LBD) were also observed. Additionally, one variant encoded *Ml*-Fd with the ER LBD inserted in the reverse orientation following Fd residue 33.

To establish whether the variants arising from LBD insertion within the Fd gene support ET in cells, we compared their growth complementation with native *Ml*-Fd and a mutant (C42A) that is unable to coordinate a 2Fe-2S metallocluster and support ET from FNR to SIR^3^. These experiments were performed in cells constitutively expressing *Zea mays* FNR and SIR and in liquid medium containing only sulfate as a sulfur source. Four out of the five sFd-ER variants presented growth complementation that is significantly higher than that observed with *Ml*-Fd-C42A in the presence of 4-HT, including sFd-30-ER, sFd-35-ER, sFd-54-ER, and sFd-55-ER (Figure 2b). This finding suggests that these fusion proteins all acquire a 2Fe-2S metallocluster and are capable of supporting ET from FNR to SIR. sFd-35-ER is related to a variant that was rationally designed in a previous study^3^. The sFd-35-ER discovered in our library has two additional residues (Ile-Asp) before the linker that connects the N-terminus of the ER LBD to *Ml*-Fd. These extra residues are added because MuA insertion of the transposon leads to a short sequence duplication (5 bp) at the insertion site ^52^.

### Evaluating ligand regulation

Our selection was performed in the presence of 4-HT to identify sFd-ER that support ET in the presence of antagonist. To determine whether any of these variants require antagonist for efficient ET in cells, we examined the effects of different 4-HT concentrations on the growth of *E. coli* EW11 coexpressing each sFd-ER variant, *Zea mays* FNR, and *Zea mays* SIR. As controls, we performed similar experiments with *Ml*-Fd, which supports ET from FNR to SIR, and *Ml*-Fd-C42A, which does not support ET from FNR to SIR^3^. In the absence of 4-HT, these experiments revealed that all of the sFd-ER variants present growth complementation that is significantly lower than that of *Ml*-Fd (Figure 2b), but not significantly higher than the C42A mutant (*P* > 0.05, using a two-tailed, independent t-test). At the lowest 4-HT concentration tested (0.781 µM), sFd-54-ER and sFd-55-ER presented complementation that was significantly higher than that observed with the C42A mutant. At the highest 4-HT concentration (50 µM), these variants exhibited complementation that was not significantly different from *Ml*-Fd (*P* > 0.05, using a two-tailed, independent t-test). Taken together, these findings show that sFd-54-ER and sFd-55-ER exhibit chemical-dependent ET. sFd-30-ER and sFd-35-ER also displayed 4-HT induced complementation. However, the level of complementation observed at each 4-HT concentration was lower than that observed with sFd-54-ER and sFd-55-ER.

The chemical-dependent complementation observed with the sFd-ER fusion proteins could arise because 4-HT affects protein expression or because this ligand binds each sFd-ER and alters ET. To evaluate whether 4-HT influences protein expression, we created constructs that express the different sFd-ER with red fluorescent protein fused at their C-terminus through a twelve amino acid linker (GGSGGSGGSGGS) and evaluated the ratio of whole cell red fluorescence in the presence and absence of 4-HT (Figure 3). As a control, we performed identical experiments with a *Ml*-Fd-RFP fusion protein. With each RFP fusion protein, this analysis revealed similar red fluorescence in the presence and absence of 4-HT (*P* > 0.05, using a two-tailed, independent t-test). These findings provide evidence that the increased complementation observed with each fusion protein in the presence of 4-HT arises because this chemical alters the ET of each sFd-ER, rather than expression.

**Figure 3.**
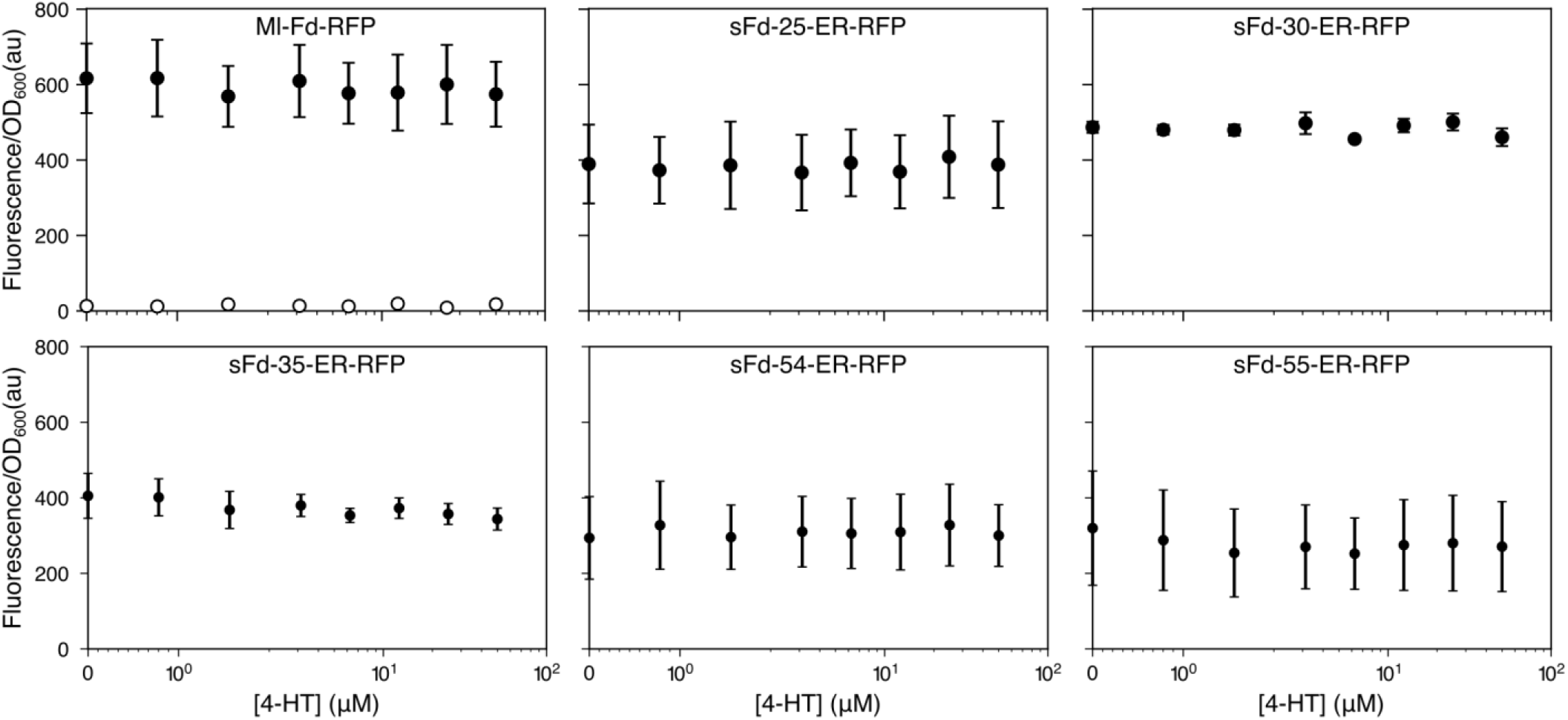
The effect of 4-HT on the fluorescence of cells expressing Fd-RFP fusions. The OD-normalized fluorescence obtained over a range of 4-HT concentrations was calculated for each Fd-RFP fusions analyzed, including *Ml*-Fd-RFP, sFd-25-ER-RFP, sFd-30-ER-RFP, sFd-35-ER-RFP, sFd-54-ER-RFP, and sFd-55-ER-RFP. As a negative control, cells containing a vector without RFP were evaluated (open circles shown with *Ml*-Fd-RFP). Circles and error bars represent the average and ±1σ, respectively, of n=3 biologically independent samples. Upon addition of 4-HT in DMSO, there was no significant increase in fluorescence compared to DMSO alone for any of the Fds tested (*P* > 0.05 using a two-tailed, independent t-test).

### In vitro measurements

In a previous study^3^, we found that recombinant sFd-35-ER contains a 2Fe-2S following purification under aerobic conditions. In this protein, domain insertion occurred outside of the *Ml*-Fd tetracysteine motif that coordinates the 2Fe-2S metallocluster^33^. To evaluate whether domain insertion within the tetracysteine motif affects the ability of a sFd-ER to acquire a 2Fe-2S cluster, we overexpressed sFd-55-ER in *E. coli* Rosetta(DE3) and purified this protein aerobically using a combination of ion exchange and size exclusion chromatography. Recombinant sFd-55-ER was brown in color following the purification protocol, like *Ml*-Fd. In addition, this protein presented a visible circular dichroism (CD) spectrum that is similar to the spectrum observed with *Ml*-Fd (Figures 4a) and other plant-type 2Fe-2S Fds^3, 53, 54^. The presence of a 2Fe-2S cluster on purified recombinant sFd-55-ER following expression shows that this protein can acquire a metallocluster in the absence of 4-HT, and it suggests that this protein functions as a chemical-dependent ET switch by regulating the conformation of the protein rather than regulating the acquisition of the redox active iron-sulfur cluster.

To investigate if domain insertion affects the reduction potential of *Ml*-Fd, we compared the electrochemical properties of sFd-55-ER with *Ml*-Fd using protein thin film voltammetry. Experiments with sFd-55-ER could not be performed at pH = 7.0 in the presence of neomycin, as in a previous study^3^, because sFd-55-ER precipitated when concentrated under these conditions. For this reason, we performed experiments under more basic conditions (pH = 8.0) in the absence of neomycin. Under this condition, the midpoint potential of *Ml*-Fd was −428 mV (Figure 4b), a value that is 92 mV more negative than that observed at pH = 7.0 previously^3^. The midpoint potential of sFd-55-ER (−418 mV) was 10 mV more positive than *Ml*-Fd (Figure 4c). Experiments performed in the presence of 4-HT revealed that the *Ml*-Fd midpoint potential was not affected by this chemical, while the midpoint potential of sFd-55-ER shifted 45 mV more negative.

**Figure 4.**
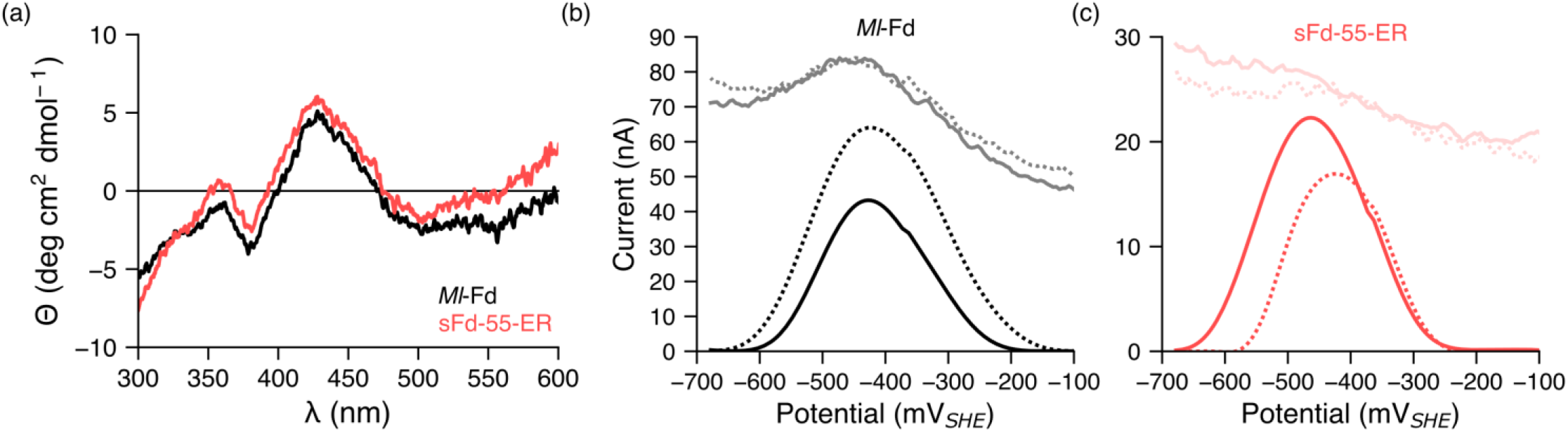
Characterization of a purified Fd-ER-LBD fusion. (**a**) Visible CD spectra of sFd-55-ER (red) and *Ml*-Fd (black) acquired under aerobic conditions at 20°C. (**b**) *Ml*-Fd and (**c**) sFd-55-ER were subjected to protein thin-film voltammetry in the absence (dashed line) and presence (solid line) of 4-HT. Baseline-subtracted data are plotted at the bottom, while raw square wave voltammetry data are shown at the top.

### Mapping domain insertion onto structure

The sFd-ER variants that exhibit chemical-dependent ET in cells arise from domain insertion at different distances from the 2Fe-2S cluster in *Ml*-Fd (Figure 5a). Within the *Ml*-Fd structure^33^, the backbone cleavage sites occur at 14.7 Å (sFd-30-ER), 18.6 Å (sFd-35-ER), 15.8 Å (sFd-54-ER), and 19.4 Å (sFd-55-ER) from the center of mass of the 2Fe-2S cluster and the alpha carbon of the residue where cleavage occurs as measured using PyMOL. We also compared the domain insertion sites with the cysteines that coordinate the iron-sulfur cluster (Figure 5b-c). This analysis shows that two of the protein variants arise from LBD insertion within the tetracysteine motif (sFd-54-ER and sFd-55-ER), and the other two arise from domain insertion between the N-terminus of *Ml*-Fd and the tetracysteine motif (sFd-30-ER and sFd-35-ER). Mapping the backbone fission sites onto the structure of the *Zea mays* Fd-SIR complex reveals that three of the variants that display chemical-dependent ET (sFd-35-ER, sFd-54-ER, and sFd-55-ER) have SIR contacting residues divided among the Fd fragments^55^. In contrast, the FNR-contacting residues observed in the *Zea mays* Fd-FNR complex are not divided among the different Fd fragments in any of the active sFd-ER that display chemical-dependent ET in cells^56^.

**Figure 5.**
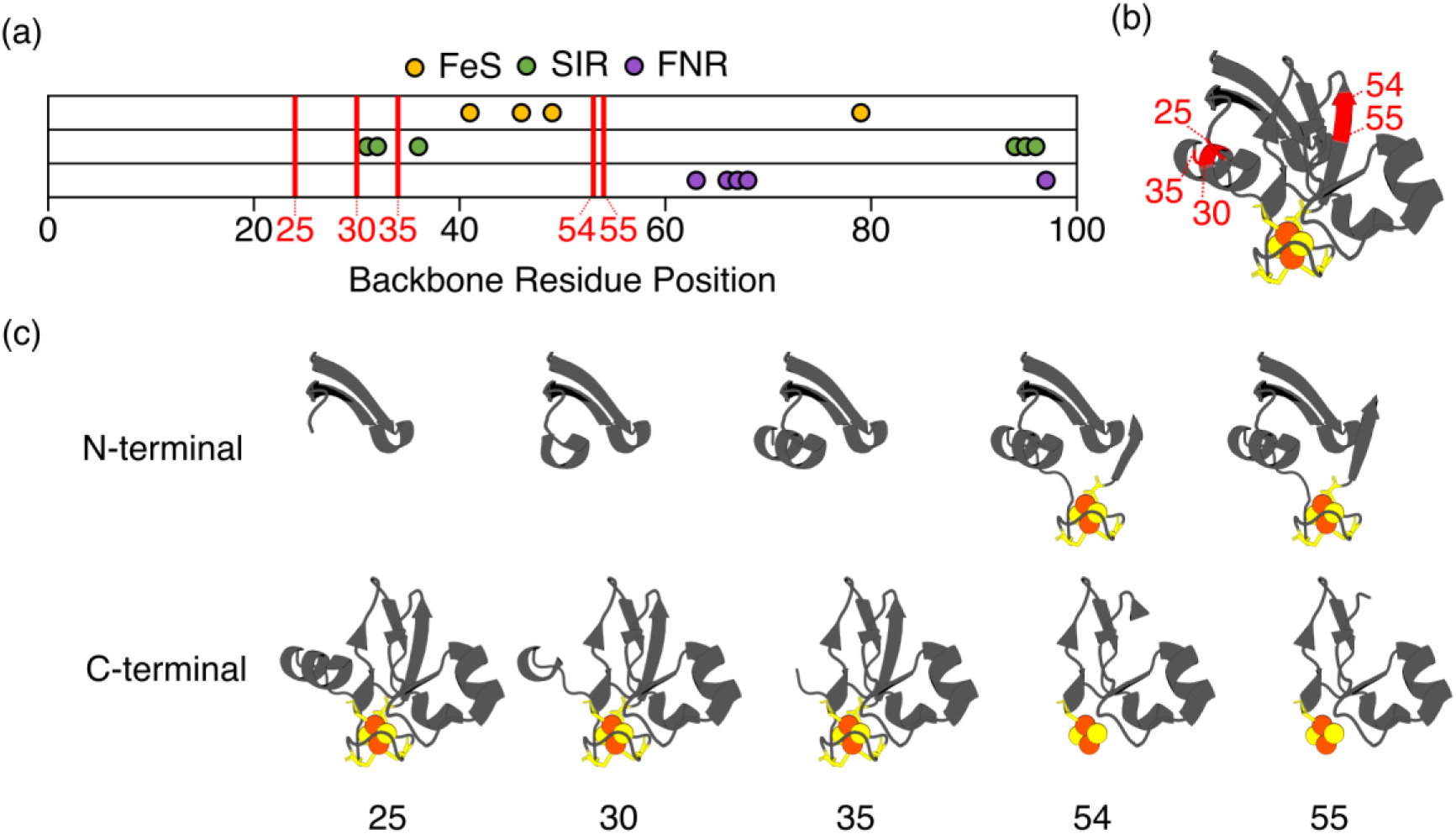
ER-LBD insertion sites mapped onto Fd structure. (**a**) A comparison of the *Ml*-Fd primary structure with the domain insertion sites (red lines) and the residues that coordinate the 2Fe-2S cluster (yellow), make residue-residue contacts SIR (green), and make contacts with FNR (purple). (**b**) A comparison of the tertiary structure of *Ml-Fd* (PDB: 1RFK) with the domain insertion sites in red. (**c**) The Fd fragments generated by domain insertion illustrate the proximity of the insertion sites and the 2Fe-2S cluster. Insertion after Leu54 or Ile55 occurs within the tetracysteine motif while the other domain insertion sites (after Gln25, Ala30, and Gly35) yield a pair of fragments with all of the cysteines on one fragment.

## Conclusion

Our combinatorial experiment identified three novel chemical-dependent ET proteins, illustrating the benefit of mining a library of topological mutants with a selection for protein design^37, 44^. Fluorescence analysis of protein expression in the presence and absence of ligand provided evidence that all three of these sFds regulate ET at the post-translational level. This ligand-induced ET could arise because of changes in: (i) iron-sulfur cluster biogenesis, (ii) iron-sulfur cluster stability, (iii) the electrochemical potential of the Fd, (iv) Fd partner binding, and/or (v) ET rates. *In vitro* studies of sFd-55-ER revealed that this protein acquires an oxygen-tolerant 2Fe-2S cluster in the absence of an ER ligand. This finding suggests that domain insertion does not regulate ET through iron-sulfur cluster biogenesis or stability. Additionally, these observations suggest that ligand does not alter the proximity of the Fd fragments fused to each end of the ER-LBD, since both fragments contain ligands required for 2Fe-2S cluster binding.

Electrochemical measurements revealed that sFd-55-ER has a midpoint potential that is more positive than *Ml*-Fd, and these measurements found that ligand binding induces a change in the midpoint potential of sFd-55-ER so that it becomes 45 mV more negative. A previous study examining the *in vitro* properties of sFd-35-ER observed a similar midpoint potential shift but in the opposite direction^3^. Our finding with sFd-55-ER suggests that ligand-induced changes in the midpoint potential of this protein is likely not responsible for the ligand-induced ET observed in cells, and it implicates changes in partner binding as the mechanism of ET regulation. To directly test this idea, additional *in vitro* studies will be required in the future that examine the effects of ligand binding on sFd-ER affinities for FNR and SIR as well as ET rates. Assays for monitoring FNR affinity for Fd interactions have been described previously^57^, and SIR affinity for Fd can be quickly measured by monitoring the effect of Fd concentration on sulfide production using a fluorescent reporter, sulfidefluor-7^58^.

Previous studies examining sequence diversity in libraries created using transposon mutagenesis have revealed large biases^52^, with the abundance of insertion sites varying over three orders of magnitude^59^. This observation suggests that our library sampling was relatively sparse, and it highlights the importance of sampling millions of vectors at each step during library construction that involves bacterial transformations. With improved sampling, the selection described herein could be coupled to deep sequencing to generate a more comprehensive profile of the locations where *Ml*-Fd tolerates domain insertion^44, 60^ and other topological mutations, like circular permutation^59^ and fission^39^. Such experiments will aid in developing design rules for creating ferredoxin switches that regulate ET for diverse bioelectronic and metabolic engineering applications.

## ACKNOWLEDGEMENTS

This research was supported by Office of Naval Research grant N00014-17-1-2639 (to J.J.S.). Additionally, JTA was supported by a Lodieska Stockbridge Vaughn Fellowship.

